# Epigenetic targeting of PGBD5-dependent DNA damage in SMARCB1-deficient sarcomas

**DOI:** 10.1101/2024.05.03.592420

**Authors:** Yaniv Kazansky, Helen S. Mueller, Daniel Cameron, Phillip Demarest, Nadia Zaffaroni, Noemi Arrighetti, Valentina Zuco, Prabhjot S. Mundi, Yasumichi Kuwahara, Romel Somwar, Rui Qu, Andrea Califano, Elisa de Stanchina, Filemon S. Dela Cruz, Andrew L. Kung, Mrinal M. Gounder, Alex Kentsis

## Abstract

Despite the potential of targeted epigenetic therapies, most cancers do not respond to current epigenetic drugs. The Polycomb repressive complex EZH2 inhibitor tazemetostat was recently approved for the treatment of *SMARCB1*-deficient epithelioid sarcomas, based on the functional antagonism between PRC2 and loss of SMARCB1. Through the analysis of tazemetostat-treated patient tumors, we recently defined key principles of their response and resistance to EZH2 epigenetic therapy. Here, using transcriptomic inference from *SMARCB1*-deficient tumor cells, we nominate the DNA damage repair kinase ATR as a target for rational combination EZH2 epigenetic therapy. We show that EZH2 inhibition promotes DNA damage in epithelioid and rhabdoid tumor cells, at least in part via its induction of the transposase-derived PGBD5. We leverage this collateral synthetic lethal dependency to target PGBD5-dependent DNA damage by inhibition of ATR but not CHK1 using elimusertib. Consequently, combined EZH2 and ATR inhibition improves therapeutic responses in diverse patient-derived epithelioid and rhabdoid tumors *in vivo*. This advances a combination epigenetic therapy based on EZH2-PGBD5 synthetic lethal dependency suitable for immediate translation to clinical trials for patients.

## Introduction

Targeted epigenetic therapies offer potential improvements over conventional cytotoxic chemotherapies through superior clinical efficacy and reduced toxicity. In cancers caused by genetic mutations of transcriptional and epigenetic regulators, specific inhibition of epigenetic effectors can directly block dysregulated gene expression by leveraging cancer-specific dependencies. One promising example of this therapeutic approach is in cancers caused by mutation of the chromatin remodeling SWI/SNF/BAF (Brg/Brahma-associated factors) complex, a ubiquitous epigenetic regulator that is mutated in at least 20% of human cancers (1). In particular, the highly lethal solid tumors caused by loss of the core BAF subunit *SMARCB1* (2–4), which include malignant rhabdoid tumors (MRT) and epithelioid sarcomas (ES), are known to be dependent on the methyltransferase EZH2, a component of the Polycomb Repressive Complex 2 (PRC2). This dependency is thought to result from epigenetic antagonism between BAF and PRC2, in which normal BAF activity evicts PRC2 from tumor suppressor loci (5, 6). In a recent clinical trial, this dependency was targeted using the EZH2 methyltransferase inhibitor tazemetostat (TAZ), leading to its FDA approval (7). Yet despite the promise of this therapeutic approach, EZH2 inhibition as a monotherapy exhibited efficacy in only a subset of patients, with most patient tumors having either primary resistance or acquiring resistance after treatment (7, 8). Thus, there is a critical need to develop improved epigenetic combination therapies, which can be achieved at least in part through improved understanding of the effects of EZH2 inhibition in epithelioid and rhabdoid tumors.

We recently leveraged functional genomics of epithelioid and rhabdoid tumors to elucidate the mechanisms of resistance to EZH2 inhibition, including frequent disruption of the RB1/E2F cell cycle control axis causing TAZ resistance in patients. This discovery led to the identification of the AURKB inhibitor barasertib combination therapy that can overcome *RB1*-mediated TAZ resistance *in vitro* and improve TAZ responses in preclinical epithelioid and rhabdoid tumor xenografts *in vivo* (8). This approach bypasses primary and acquired RB1/E2F-mediated checkpoint defects to maintain TAZ-induced reprogramming of oncogenic gene expression. However, RB1/E2F pathway alterations were observed in less than half of epithelioid and rhabdoid tumors, raising the question of additional combinations that may achieve effective epigenetic therapy for patients, including those with primary or acquired TAZ resistance.

Epithelioid and rhabdoid tumors occur due to characteristic biallelic deletions and inactivating mutations of *SMARCB1*, either as a result of germline rhabdoid tumor predisposition syndrome, or due to somatic mutations (2, 9, 10). For rhabdoid tumors, differences in the types of mutations, e.g. large versus small chromosomal deletions or missense and nonsense mutations of *SMARCB1* correspond with distinct molecular subtypes and clinical features (11). Genome sequencing studies have identified distinct sequence-specific mutational features in rhabdoid tumors, including sequence-specific deletions of *SMARCB1* and recurrent mutations of other genes, associated at least in part with the mutagenic activity of the domesticated transposase-derived Piggybac Transposable Element Derived 5 (PGBD5) (9).

PGBD5 is among the most evolutionarily conserved domesticated transposase-derived genes in vertebrates and can mediate sequence-specific DNA rearrangements dependent on its putative nuclease activity (9, 12). In particular, Pgbd5 promotes sequence-specific somatic mutagenesis and tumor development in mouse models of medulloblastoma, one of the most common childhood brain tumors (13). PGBD5-expressing cells, including rhabdoid tumor cells, require a distinct form of non-homologous end-joining (NHEJ) DNA repair, leading to their hypersensitivity to inhibition of DNA repair signaling, specifically ATR inhibition (14). We have now found that TAZ inhibition of EZH2 can increase PGBD5 expression in epithelioid and rhabdoid tumor cells, potentiating PGBD5-dependent DNA damage, and conferring synergy with the ATR-selective DNA repair inhibitor elimusertib. Accordingly, combined treatment of patient-derived rhabdoid and epithelioid tumors engrafted into immunodeficient mice, including from patients with multiply relapsed metastatic disease, demonstrates synergistic activity *in vivo*.

## Results

To define potential therapeutic targets for TAZ combination therapy in *SMARCB1*-deficient tumors, we leveraged a recently developed method for transcriptomic inference of oncogenic protein activity, termed metaVIPER (Virtual Inference of Protein activity by Enriched Regulon), as applied to gene expression profiles of 68 patient rhabdoid tumors (15–17). We used the OncoTarget annotation of high-affinity pharmacologic inhibitors of putative master gene expression regulators (MRs) to order them by their mean activity scores (**Figure 1A**). As expected, EZH2 was among the most activated MR proteins in rhabdoid tumors, as was the previously identified TAZ combination therapy target, AURKB (**Figure 1A**). This analysis also prioritized CDK4, CDK2, and AURKA (**Figure 1A**), consistent with the previously established activity of cell cycle checkpoints downstream of G1/S in rhabdoid tumors (8), validating this ranking approach.

**Figure 1:**
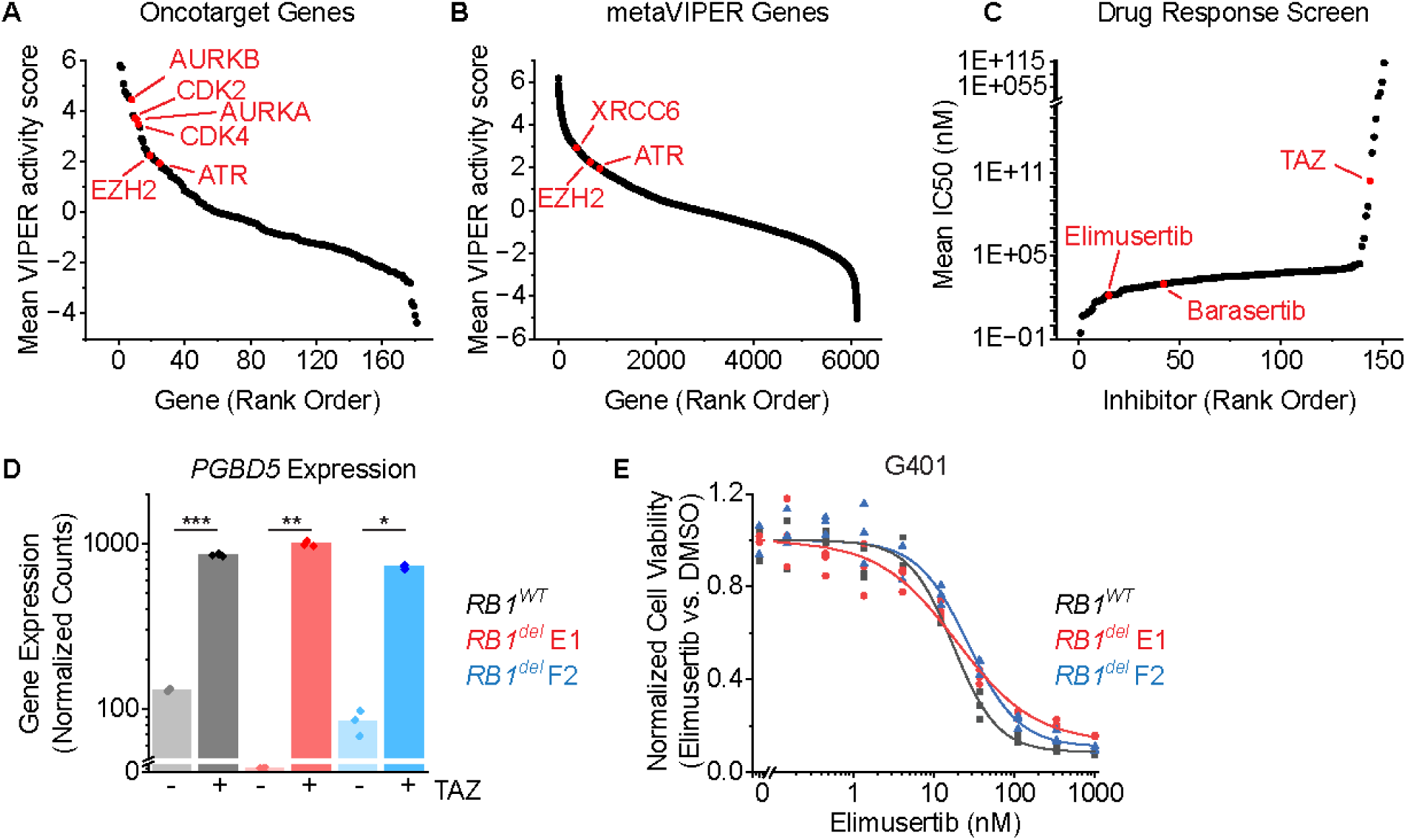
ATR inhibition is a compelling therapeutic target and overcomes TAZ resistance in rhabdoid tumors. (**A-B**) metaVIPER analysis of rhabdoid tumor transcriptomes for proteins within OncoTarget (**A**) and the complete metaVIPER protein set (**B**). (**C**) Inhibitors rank-ordered by IC50 from Kuster et al. (**D**) DESeq2-normalized read counts of cells treated with 10 µM TAZ versus equivalent volume of DMSO for 11 days. n=3 biological replicates per condition. \**p* = 1.6E-6, \*\**p* = 1.3E-6, \*\*\**p* = 1.2E-7 by two-sided Student’s t-test. (**E**) G401 cells treated with elimusertib for 4 days.

In addition to these known dependencies, this analysis also identified the NHEJ DNA repair structural factor XRCC6 (Ku70), and its mediator kinase ATR among the most activated MR genes among all 6,000 MR proteins in metaVIPER (**Figure 1B**), both of which have been shown to be required for the survival of cells expressing active PGBD5 in prior studies (14). Indeed, XRCC6 ranked higher in this analysis than the well-validated target EZH2 (**Figure 1B**).

To ascertain the robustness of this prediction, we next analyzed an independent dataset from Kuster and colleagues (18), which assessed the responses of sarcoma cell lines to a panel of 151 pharmacologic inhibitors, including G401 and A204 rhabdoid tumor and VA-ES-BJ epithelioid sarcoma cell lines. We observed that the ATR-selective kinase inhibitor elimusertib was among the most active drugs against epithelioid and rhabdoid tumor cells, even more active than both TAZ and barasertib (**Figure 1C**). Thus, *SMARCB1*-deficient epithelioid and rhabdoid tumors are highly sensitive to ATR inhibition (14).

Since both modulation of gene expression by TAZ and DNA repair by ATR inhibitor elimusertib appears to have prominent activity in rhabdoid tumor cells, we inquired whether TAZ may regulate PGBD5 itself. We found that in addition to the expected upregulation of Polycomb gene sets, EZH2 inhibition in G401 rhabdoid tumor cells with TAZ for 11 days also caused a significant increase in the expression of *PGBD5* (mean fold-increase of 6.6 and Student’s t-test *p* = 1.2E-7; **Figure 1D**). Using CRISPR gene editing, we generated isogenic *RB1* wild-type and *RB1^del^*mutant G401 cells and confirmed correct biallelic *RB1* inactivating mutations and consequent loss of RB1 protein expression, using the *AAVS1* safe harbor as a negative control (8). In prior work, we found that defects in the RB1/E2F axis, including mutations of *RB1*, cause TAZ resistance (8). Consistent with the independence of this TAZ resistance mechanism from ATR inhibitor susceptibility, two independent G401 *RB1^del^* mutant cell lines also exhibited significant TAZ-mediated induction of *PGBD5* expression (mean fold-increase of 333 and 8.6 and t-test *p* = 1.3E-6 and 1.6E-6 for E1 and F2 clones, respectively; **Figure 1D**). Additionally, both *RB1^WT^*and *RB1^del^* mutant G401 cells exhibited nanomolar susceptibility to the ATR-selective inhibitor elimusertib (half-maximal effective concentration of 18 ± 1.6 nM, 19 ± 3.8 nM, and 27 ± 3.2 nM for *RB1^WT^*, *RB1^del^* E1, and F2 clones, respectively; **Figure 1E**).

Elimusertib is currently undergoing clinical trials in patients with refractory or relapsed solid tumors, including patients with PGBD5-expressing tumors such as MRT and ES (Clinical Trials Identifier NCT05071209). We therefore investigated the activity of combination treatment with TAZ and elimusertib, reasoning that TAZ-induced upregulation of *PGBD5* expression may potentiate the anti-tumor effects of ATR inhibition. We observed greater antitumor effects from the combination of TAZ and elimusertib than either drug alone against diverse MRT and ES cell lines, including TAZ-resistant ES1 and VAESBJ cells (mean fold-decrease in normalized cell viability of 0.64, 0.63, 0.78, 0.84, and 0.84 compared with most effective monotherapy for TTC642, KP-MRT-NS, KP-MRT-RY, G401 and ES1 respectively, t-test 1.0E-3, 3.0E-5, 3.3E-3, 2.6E-4, and 6.1E-5;**Figure 2A**). Strikingly, the combination of TAZ and elimusertib was highly synergistic in G401 rhabdoid and ES1 epithelioid sarcomas cells (ZIP synergy = 3.4 and 2.0 and *p* = 9.8E-1 and 7.3E-1, respectively; **Figure 2B-C**).

**Figure 2:**
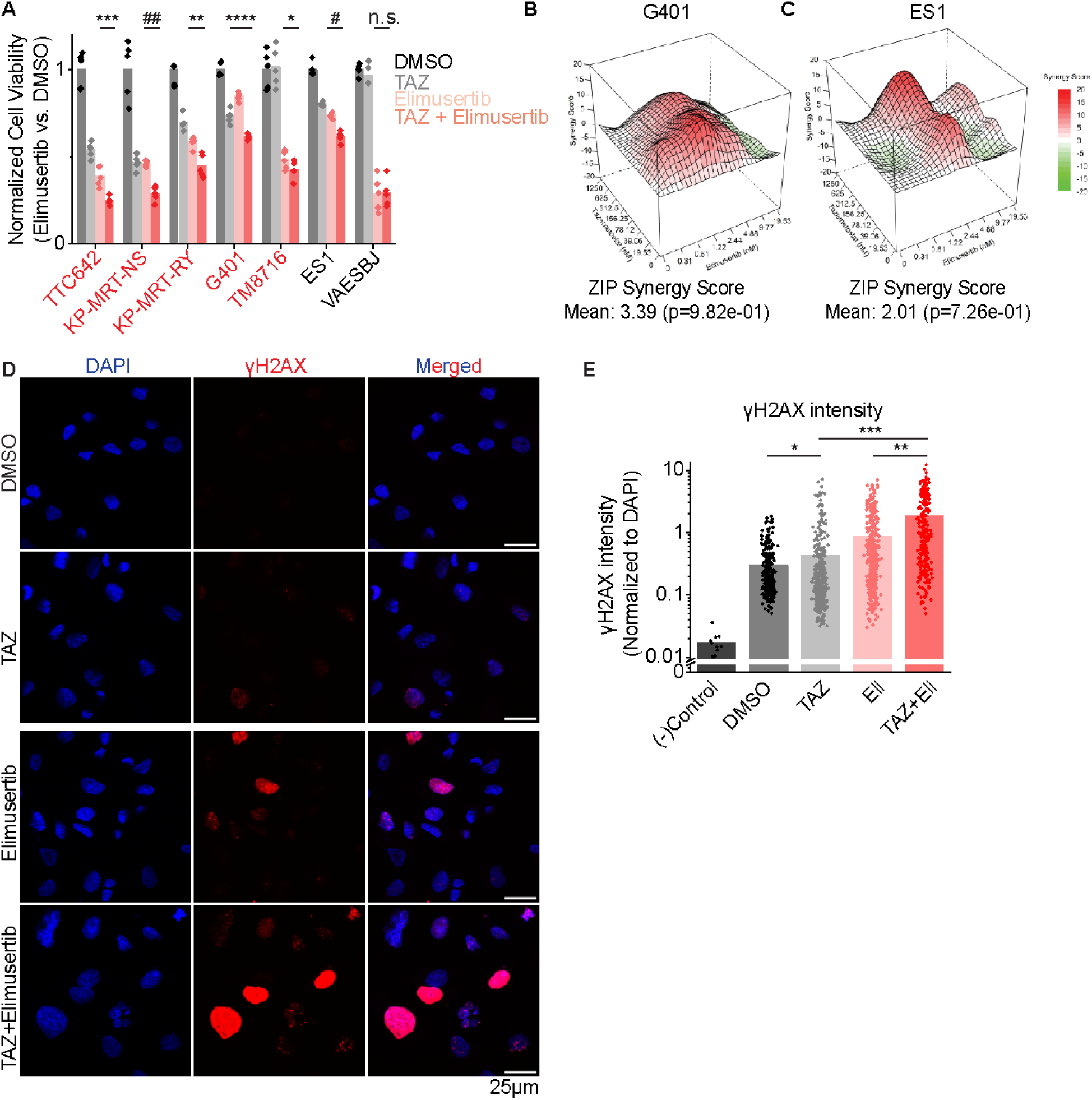
Combination EZH2 and ATR inhibition improves response in vitro. (**A**) Panel of MRT and ES cell lines ordered left to right by decreasing response to TAZ monotherapy. Cells were treated with the indicated monotherapy or combination for 11 days. Drug concentrations used were: TAZ: 200 nM, elimusertib: 8 nM. We selected an elimusertib dose below its monotherapy IC_50_ (for G401 cells) in order to visualize any additive effects upon combination with TAZ. \**p* = 0.14, \*\**p* = 3.3E-3, \*\*\**p* = 1.0E-3, \*\*\*\**p* = 2.6E-4, #*p* = 6.1E-5, ##*p* = 3.0E-5 by two-sided Student’s t-test. All comparisons refer to TAZ + elimusertib vs. elimusertib conditions, except for G401 cells, in which comparison is for TAZ + elimusertib vs. TAZ. n = 5 replicates per condition. (**B-C**) Synergy plots for combination treatment with TAZ (left axis) and elimusertib (right axis) for (**B**) G401 and (**C**) ES1 cells. Cells were treated at the indicated doses for 9 days and analyzed for synergy using the Zero Interaction Potency (ZIP) model. (**D**) Representative images of G401 cells treated with the indicated treatment for 7 days. Elimusertib was added on Day 5. Doses used were 500 mM TAZ and 100 nM elimusertib. (**E**) Quantification of γH2AX fluorescence relative to DAPI fluorescence. \**p* = 9.3E-3, \*\**p* = 6.2E-14, \*\*\**p* = 1.5E-29 by two-sided Student’s t-test. n = 332 nuclei for DMSO, 404 for TAZ, 400 for elimusertib, 257 for TAZ + elimusertib.

If the improved anti-tumor activity of TAZ and elimusertib is due to the induction of PGBD5-dependent DNA damage, then this treatment combination should be associated with the induction of DNA damage repair signaling. To test this prediction, we used confocal immunofluorescence microscopy to quantify γH2AX as a specific marker of DNA damage (19). In agreement with prior studies (14), vehicle-treated G401 cells showed measurable γH2AX staining associated with baseline *PGBD5* expression (**Figure 2D-E**). Consistent with TAZ-mediated induction of *PGBD5* expression (**Figure 1D**), we found that TAZ treatment alone significantly increased nuclear γH2AX (mean normalized intensity = 0.42 versus 0.29 for TAZ and DMSO, respectively; *t*-test *p* = 9.3E-3; **Figures 2D-E**). The combination of TAZ and elimusertib induced a significant increase in γH2AX as compared to either drug alone (mean normalized intensity = 1.8 for the combination versus 0.42 and 0.85 for TAZ and elimusertib, respectively; *t*-test *p* = 6.2E-14 and 1.5E-29 for combination versus elimusertib and TAZ, respectively; **Figures 2D-E**).

Recently, EZH2 suppression was shown to induce replication stress through upregulation of MYCN expression in T-ALL cells, which in turn sensitized cells to inhibition of CHK1 (19), a downstream mediator of ATR signaling (20). To test this possibility, we measured *MYCN* expression in TAZ-treated G401 rhabdoid tumor cells, which was significantly increased (**Figure 3A**). However, this induction of *MYCN* gene expression was not associated with the accumulation of MYCN protein, as measured by Western blotting with *MYCN*-amplified IMR5 neuroblastoma cells as a positive control (**Figure 3B**). Consistent with this, EZH2 inhibition either alone or in combination with CHK1 inhibition did not induce apparent replication stress, as measured by RPA phosphorylation, using the treatment of cells with the DNA topoisomerase inhibitor camptothecin as positive control for replication stress. This was despite the effective inhibition of CHK1 auto-phosphorylation by the CHK1-selective SRA737 inhibitor (**Figure 3C**). Concordantly, the CHK1 inhibitor SRA737 showed poor activity against G401 cells, regardless of *RB1* status (**Figure 3D**). Thus, rhabdoid tumor cells exhibit a specific dependency on ATR-dependent but CHK1-independent DNA damage repair signaling.

**Figure 3:**
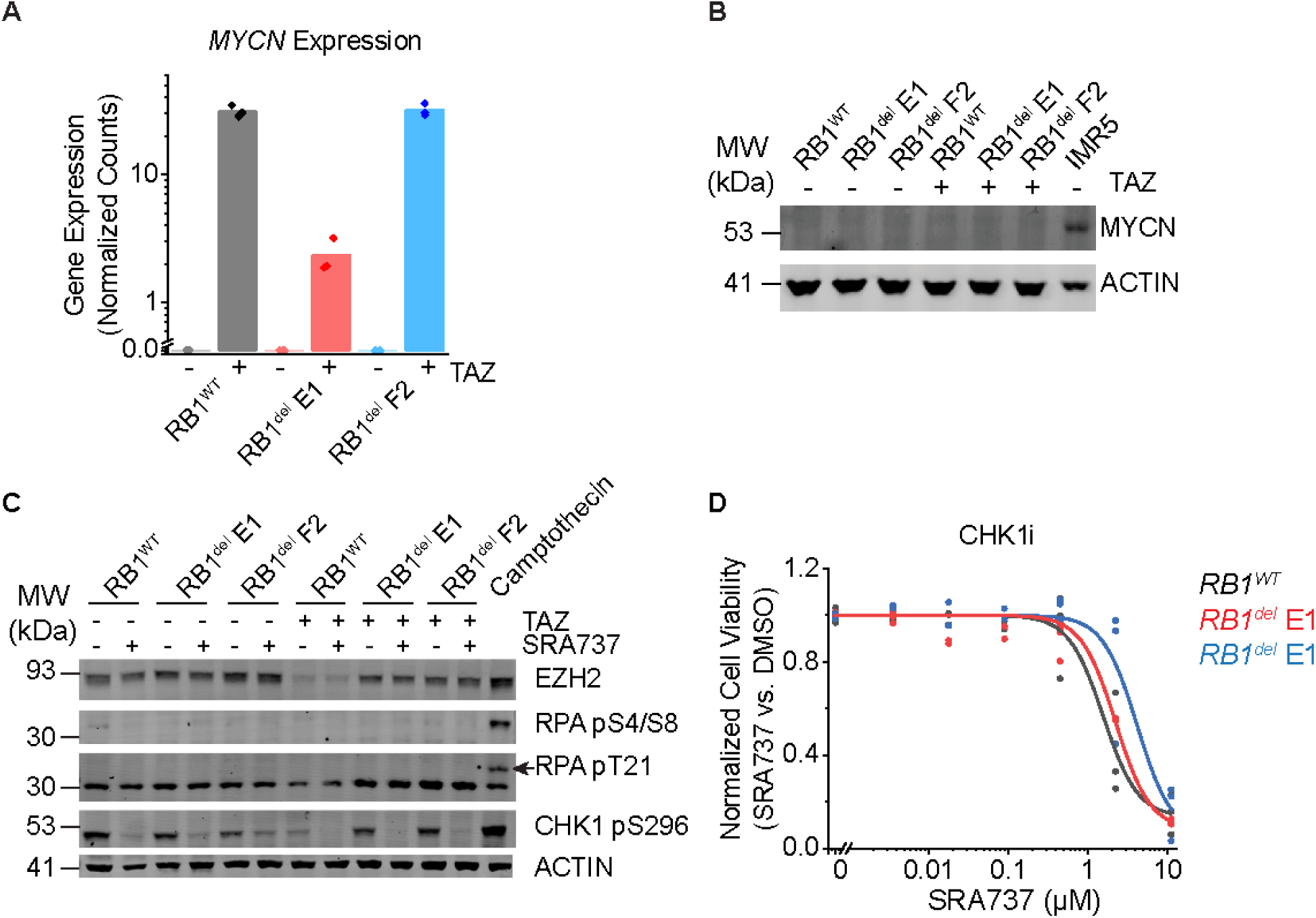
CHK1 inhibition does not induce replication stress or synergize with TAZ. (**A**) DESeq2-normalized read counts of cells treated with 10 µM TAZ versus equivalent volume of DMSO for 11 days. n=3 biological replicates per condition. (**B**) Dose-response curves of G401 cells treated with the CHK1 inhibitor SRA737 for 9 days. (**C**) Western blot assaying replication stress as measured by RPA phosphorylation at S4/8 and T21. Camptothecin treatment (1.5 µM) for 2 h was used as a positive control for replication stress. Autophosphorylation of CHK1 at S296 was used to confirm CHK1 inhibition. Cells were pre-treated with 10 µM TAZ or DMSO for 9 days. Cells were then split and additionally treated with SRA737 (3 µM) or equivalent volume of DMSO for 2 days. (**D**) Cells treated with 10 µM TAZ or DMSO for 11 days do not express MYCN protein. *MYCN*-amplified neuroblastoma cell line IMR5 was used as a positive control for MYCN expression.

TAZ-mediated induction of *PGBD5* and DNA damage would indicate that this form of DNA damage should be dependent on *PGBD5* expression. To test this prediction, we engineered short hairpin RNA-mediated depletion of *PGBD5* in G401 cells using lentiviral transduction of two independent *PGBD5*-specific shRNAs (shPGBD5), as compared to a control GFP-targeting shRNA (shGFP). We confirmed that shPGBD5-expressing cells were significantly depleted of PGBD5 as compared to control shGFP cells (mean fold-depletion = 0.59 and 0.54; t-test *p* = 4.7E-3 and 1.2E-3, respectively; **Figure 4A**). We then measured DNA damage using quantitative confocal immunofluorescence microscopy of γH2AX, and found that while the combination of TAZ and elimusertib induced DNA damage in shPGBD5 cells, this effect was significantly reduced compared to control shGFP cells (mean fold-reduction = 0.34 and 0.39; t-test *p* = 3.9E-8 and 4.4E-8 for shGFP versus shPGBD5-1 and shPGBD5-3, respectively; **Figure 4B-C**). This PGBD5-dependent reduction in DNA damage was observed both when examining all nuclei with γH2AX staining (**Figure 4B**), as well as only nuclei with punctate γH2AX staining, corresponding to more localized DNA damage (mean fold-reduction = 0.59 and 0.61; t-test *p* = 2.4E-5 and 2.7E-5 for shGFP versus shPGBD5-1 and shPGBD5-3, respectively;**Figure 4D**), as opposed to pan-nuclear γH2AX staining due to genome-wide unrepaired DNA damage and cellular apoptosis (mean fold-reduction = 0.31 and 0.55; t-test *p* = 9.0E-4 and 1.6E-2 for shGFP versus shPGBD5-1 and shPGBD5-3, respectively; **Figure 4E**). Thus, PGBD5 is at least in part necessary for TAZ-mediated induction of DNA damage and its potentiation by the combination with elimusertib.

**Figure 4:**
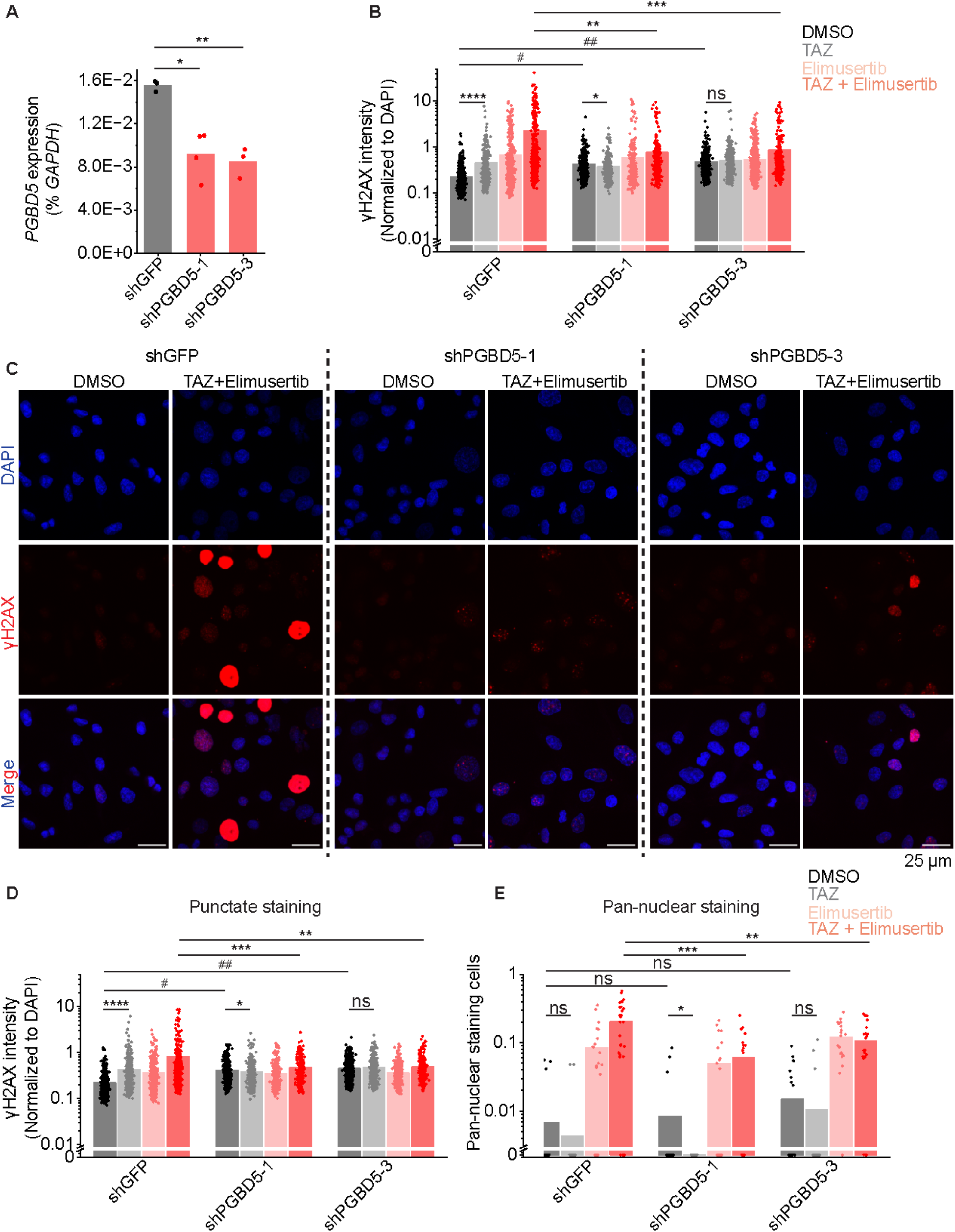
TAZ-induced DNA damage is PGBD5-dependent. (**A**) RT-qPCR showing *PGBD5* expression vs. *GAPDH* in G401 cells with the indicated shRNA. \**p* = 4.7E-3, \*\**p* = 1.2E-3. (**B**) Quantification of γH2AX fluorescence relative to DAPI fluorescence in all nuclei. \**p* = 8.6E-3, \*\**p* = 3.9E-8, \*\*\**p* = 4.4E-8, \*\*\*\**p* = 1.1E-12, # *p* = 4.9E-34, ## *p* = 4.0E-47 by two-sided Student’s t-test. (**C**) Representative images of cells quantified in (B). (**D**) Quantification of γH2AX fluorescence relative to DAPI fluorescence in nuclei with punctate γH2AX staining. \**p* = 3.6E-2, \*\**p* = 2.7E-5, \*\*\**p* = 2.4E-5, \*\*\*\**p* = 1.1E-15, #*p* = 5.6E-51, ##*p* = 1.7E-81. For B-C, n for shGFP cells is 539 for DMSO, 321 for TAZ, 491 for elimusertib, 418 for TAZ + elimusertib. n for shPGBD5-1 is 497 for DMSO, 390 for TAZ, 354 for elimusertib, 277 for TAZ + elimusertib. n for shPGBD5-3 is 703 for DMSO, 418 for TAZ, 554 for elimusertib, 317 for TAZ + elimusertib. (**E**) Proportion of nuclei with pan-nuclear γH2AX staining per field. Each dot represents one field. \**p* = 9.2E-2, \*\**p* = 1.6E-2, \*\*\**p* = 9.0E-4 n for shGFP cells is 22 fields for DMSO, 22 for TAZ, 22 for elimusertib, 28 for TAZ + elimusertib. n for shPGBD5-1 is 22 for DMSO, 22 for TAZ, 22 for elimusertib, 20 for TAZ + elimusertib. n for shPGBD5-3 is 22 for DMSO, 22 for TAZ, 20 for elimusertib, 22 for TAZ + elimusertib.

This nominates therapeutic targeting of the EZH2-PGBD5 synthetic lethal dependency as an improved combination strategy for epithelioid and rhabdoid tumors. To test this idea, we assembled a phase 2-like cohort of diverse MRT and ES tumors derived from patients with relapsed and metastatic disease, including tumors with numerous additional acquired mutations (**Supplementary Table S1**). We engrafted these tumors into immunodeficient *NOD-scid IL2Rgamma^null^* mice, and randomized tumor-bearing animals to treatment with TAZ or elimusertib or the combination of both (**Figure 5A**). The combination of TAZ and elimusertib exceeded the effect of treatment with either drug alone when assessed by tumor growth measurements (Vardi *U*-test *p* = 2.33E-2 and 0.20 for combination versus elimusertib or TAZ, respectively; **Figure 5A**) and significantly extended tumor-free survival from 51 days (95% CI = 42-60 days) for elimusertib and 68 days (95% CI = 53-82 days) for TAZ to 100 days (95% CI = 76-124 days) for the combination (log-rank test *p* = 5.8E-4 and 3.8E-2 for combination versus elimusertib or TAZ, respectively; **Figure 5B**).

**Figure 5:**
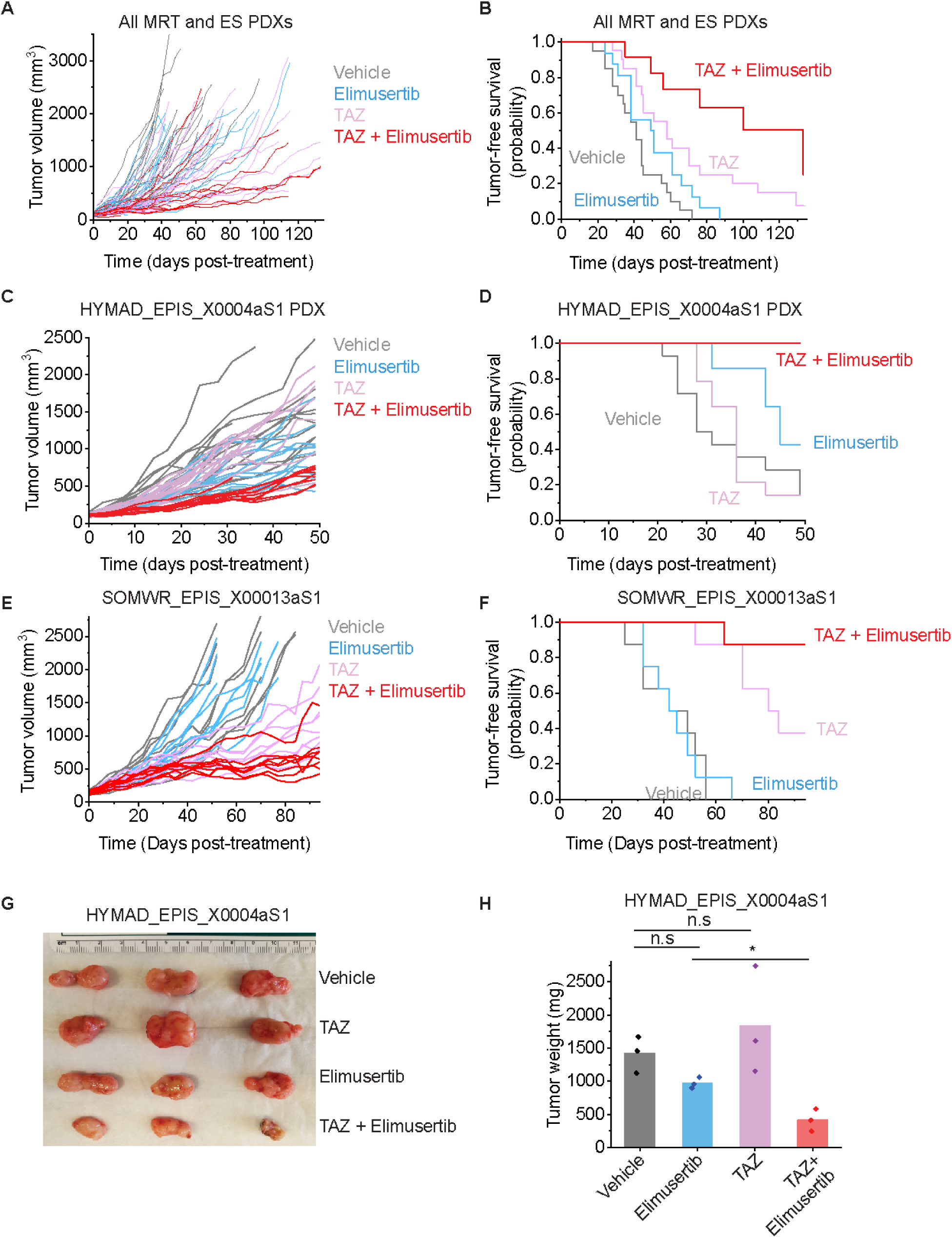
TAZ plus elimusertib improves therapeutic response in vivo. (**A**) Tumor growth curves for 5 mouse PDXs treated with the indicated drug regimen. n = 20 mice for vehicle and elimusertib-treated groups, n = 21 for TAZ and TAZ + elimusertib-treated groups. Vardi *U*-test *p* = 2.3E-2 and 0.20 for combination vs. elimusertib or TAZ, respectively. (**B**) Kaplan-Meier curves showing tumor-free survival (defined as tumor volume ≤ 1,000 mm^3^) for the PDXs in panel C. Mean survival is 51 days (95% CI: 42-60 days) for elimusertib, 68 days (95% CI: 53-82 days) for TAZ to 100 days (95% CI: 74-124 days) for the combination. Log-rank test *p* = 5.8E-4 and 3.8E-2 for combination versus elimusertib or TAZ, respectively. (**C**) Tumor growth curves for the HYMAD_EPIS_X0004aS1 PDX model treated with the indicated drug regimen. Vardi *U*-test *p* = 2.0E-4 for combination versus elimusertib or TAZ. n = 14 mice per treatment group. (**D**) Kaplan-Meier curves showing tumor-free survival (defined as tumor volume ≤ 1,000 mm^3^) for the PDXs in panel C. Log-rank test *p* = 6.2E-3 and 6.3E-5 for combination versus elimusertib or TAZ, respectively. (**E**) Tumor growth curves for the SOMWR_EPIS_X00013aS1 PDX model. Vardi *U*-test *p* = 1.0E-3 and 6.0E-2 for combination versus elimusertib and TAZ, respectively. *n* = 8 mice for all groups. (**F**) Kaplan-Meier curves showing tumor-free survival (defined as tumor volume ≤ 1,000 mm^3^) for the SOMWR_EPIS_X00013aS1 PDX model; Mean survival is 90 days (95% CI: 83-97 days) for combination versus 80 days (95% CI: 70-90 days) for TAZ and 45 days (95% CI: 37-52 days) for elimusertib. Log-rank test *p* = 9.5E-5 and 6.0E-2 for combination versus elimusertib and TAZ, respectively. (**G**) Image of representative tumors extracted from mice in C-D and on Day 52 of treatment (**H**) their corresponding weights. \**p* = 6.7E-3 by two-sided Student’s t-test.

This activity was most pronounced for the HYMAD_EPIS_X0004aS1 and SOMWR_EPIS_X00013aS1 PDX models (**Figures 5C-H**), despite the former exhibiting a relatively poor response to TAZ monotherapy, when assessed by tumor growth measurements (Vardi *U*-test *p* = 2.0E-4 for combination versus elimusertib or TAZ for HYMAD_EPIS_X0003aS1; **Figure 5C**; *p* = 1.0E-3 and 6.0E-2 for combination versus elimusertib and TAZ, respectively for SOMWR_EPIS_X00013aS1; **Figure 5E**) and tumor-free survival (log-rank test *p* = 6.2E-3 and 6.3E-5 for combination versus elimusertib or TAZ, respectively for HYMAD_EPIS_X0003aS1; **Figure 5D**; *p* = 9.5E-5 and 6.0E-2 for combination versus elimusertib or TAZ, respectively for SOMWR_EPIS_X00013aS1; **Figure 5F**). This effect leverages the EZH2-PGBD5 collateral synthetic lethal dependency, targeting PGBD5-dependent DNA damage to improve TAZ clinical response and overcome resistance in epithelioid and rhabdoid tumors.

## Discussion

Prior studies showed that inhibition of EZH2 methyltransferase activity using tazemetostat is insufficient to induce durable tumor regressions in most patients with epithelioid and rhabdoid tumors (8). Our recent functional genetic studies of patient tumors before and after clinical TAZ therapy led to a specific model of effective epigenetic therapy, including rational combinations to overcome RB1/E2F pathway defects that were observed in 43% of tumors with primary or acquired TAZ resistance (8). Here, we further advanced this approach by identifying a collateral synthetic lethal dependency between EZH2 and PGBD5 in rhabdoid and epithelioid sarcomas that confers a susceptibility to combined epigenetic therapy using EZH2 and ATR inhibition. We found that EZH2 inhibition can upregulate the transposase-derived PGBD5 nuclease, which induces DNA damage, requiring ATR but not CHK1-mediated DNA damage repair signaling in *SMARCB1*-deficient rhabdoid and epithelioid tumors regardless of their *RB1* mutational status. As a result, combined EZH2 and ATR inhibition exerts synergistic anti-tumor effects, as measured by DNA damage induction *in vitro* and tumor growth reduction and improvements in tumor-free survival *in vivo*.

The combination of EZH2 and ATR inhibition offers both a promising therapeutic approach that can be rapidly translated to clinical trials for patients with rhabdoid and epithelioid sarcomas, and a compelling example of drug-induced synthetic lethality. Originally described as genetic interactions (21), synthetic lethal targeting has proven to be a powerful therapeutic approach for the treatment of cancers. For example, breast and ovarian carcinomas with *BRCA1/2* mutations exhibit increased dependence on poly(adenosine diphosphate-ribose) polymerase (PARP)-mediated DNA repair, conferring a susceptibility to PARP1/2 inhibitors such as olaparib (22, 23). Synthetic dependencies in DNA damage signaling, chromatin remodeling, and metabolic functions have recently been defined to develop improved targeted therapies (24–28). In particular, acquired defects in DNA repair in tumor cells, such as inactivating mutations or functional defects in ATM signaling, can confer susceptibility to ATR inhibition due to the specific requirements of concurrent ATM and ATR signaling for efficient NHEJ DNA repair and tumor cell survival (28–32).

ATR-selective inhibitors also synergize with genotoxic chemotherapies, such as DNA cross-linking platinum drugs, due to the specific requirements of ATR-dependent DNA repair during DNA replication (28). However, this approach has limited therapeutic efficacy due to its effects on healthy cells. Thus, effective targeting of DNA damage repair must leverage tumor-specific mutational and repair processes, providing a rationale for combining epigenetic and DNA repair inhibitors. In the case of rhabdoid and epithelioid sarcomas with *SMARCB1* deficiency, the epigenetic antagonism between the chromatin remodeling activities of BAF and PRC2 contributes to the dependency of tumor cells on EZH2 activity (5), whose inhibition promotes the expression of tumor suppressors otherwise repressed by PRC2 (8, 33). This epigenetic reprogramming also appears to increase the expression of PGBD5, with the associated DNA damage and requirement for its ATR-dependent repair. It remains to be defined whether this effect is due to direct repression through PRC2-mediated histone 3 lysine 27 trimethylation of the *PGBD5* locus and/or indirect transcriptional or post-transcriptional regulation of *PGBD5* expression.

Though the cell of origin of rhabdoid and epithelioid sarcomas is currently unknown, these tumors exhibit epigenetic and transcriptional features of neuronal and neural crest cells (34–36). EZH2 is known to regulate neuronal differentiation (37–40), and therefore it will also be important to determine whether EZH2 inhibition of other PGBD5-expressing tumors, such as neuroblastomas, medulloblastomas, and Ewing and other fusion sarcomas, all of which also share features of neuronal lineages, can also induce PGBD5-dependent DNA damage, thereby conferring a susceptibility to collateral synthetic lethal targeting with ATR inhibitors. Importantly, susceptibility to ATR inhibition can also result from PGBD5-independent sources of intrinsic DNA damage, such as alternative lengthening of telomeres (ALT) (41), replication stress due to transcriptional-replication interference (42–44), and functional ATM defects (30, 45), all of which may occur concurrently in tumor cells undergoing PGBD5-dependent DNA damage. Similarly, EZH2 may also contribute to other mechanisms of DNA damage repair (46–53). Thus, future biochemical and genetic studies will be needed to define specific mechanisms of EZH2 and ATR-dependent DNA damage repair signaling in tumor and healthy cells to identify improved targets to develop exclusively tumor-selective therapies. This direction is particularly compelling because PGBD5-dependent DNA damage repair appears to require ATR but not CHK1 kinase signaling in epithelioid and rhabdoid tumor cells, in contrast to ATR/CHK1-dependent canonical signaling observed during replication stress in healthy tissues.

Further definition of the EZH2-PGBD5 collateral synthetic lethal mechanisms should also aid in therapy stratification of combined EZH2 and ATR inhibition. For example, apparent variation in response to the tazemetostat-elimusertib combination between patient-derived epithelioid and rhabdoid tumors may result from biological differences in DNA damage repair signaling, PGBD5 activity, or other sources of intrinsic DNA damage, which in turn may be associated with the recently described distinct molecular subtypes of ES and MRT tumors (54). In addition, both EZH2 and ATR inhibition can be immunogenic (8, 55–57), and future studies will be needed to define immunologic effects of this combination therapy that may contribute to therapeutic effects in patients. In all, this work emphasizes how improved understanding of collateral dependencies of intrinsic mutators and epigenetic dysregulation responsible for causing childhood and young-onset cancers can be leveraged for rational combination therapies.

## Supporting information

Uncropped Blots

## Acknowledgements

This work is dedicated to Maggie Schmidt and her family, and to the many other patients and their advocates who inspire and support our research. We thank Richard Koche, Nicholas Socci, and Mithat Gönen for technical advice, Marc Ladanyi for patient-derived xenografts, members of our labs for critical advice and manuscript comments, and Epizyme (now Ipsen) and Bayer for supplying tazemetostat and elimusertib, respectively. This work was supported by the MSK Integrated Genomics Operation Core, Anti-Tumor Assessment Core, Bioinformatics Core, Molecular Diagnostics Service and the Department of Pathology, the Marie-Josée and Henry R. Kravis Center for Molecular Oncology, NIH R01 CA214812, P30 CA08748, T32 GM007739, Burroughs Wellcome Fund, Rita Allen Foundation, Pershing Square Sohn Cancer Research Alliance and the G. Harold and Leila Y. Mathers Foundation, Cycle for Survival, MSK Sarcoma Center, the Starr Cancer Consortium, and Maggie’s Mission. Yaniv Kazansky was supported by a Medical Scientist Training Program grant from the National Institute of General Medical Sciences of the National Institutes of Health under award number T32 GM007739 to the Weill Cornell/Rockefeller/Sloan Kettering Tri-Institutional MD-PhD Program. Alex Kentsis is a Scholar of the Leukemia & Lymphoma Society. This work was also supported by the NCI Office of Cancer Target Discovery and Development (CTD^2^) award U01 CA 272610, the NCI Outstanding Investigator award R35 CA 197745, and the NIH Shared Instrumentation Grants S10 OD012351, S10 OD021764 and S10OD032433, all to Andrea Califano.

## Methods

### Cell culture

G401, TTC642, VAESBJ, and TM8716 cell lines were obtained from the American Type Culture Collection. The ES1 cell line was generated and kindly provided by Nadia Zaffaroni. Rhabdoid tumor cell lines KP-MRT-NS and KP-MRT-RY were kindly provided by Yasumichi Kuwahara and Hajime Hosoi. The identity of all cell lines was verified by STR analysis. Absence of *Mycoplasma* contamination was determined at every plating using the MycoAlert kit according to manufacturer’s instructions (Lonza). Cell lines were cultured in 5% CO_2_ in a humidified atmosphere in 37°C. All media were obtained from Corning and supplemented with 10% fetal bovine serum (FBS), 1% L-glutamine, and 100 U/mL penicillin and 100 µg/mL streptomycin (Gibco). G401, ES1, and VAESBJ cells were cultured in Dulbecco’s Modified Eagle Medium (DMEM). TTC642, TM8716, KP-MRT-NS, and KP-MRT-RY cells were cultured in Roswell Park Memorial Institute (RPMI) medium. All experiments were performed using cell lines kept in culture for no more than 10 passages. Generation of G401 with *RB1* deletion was described previously (8) using the gRNA sequence in Table 1 below.

**Table 1:**
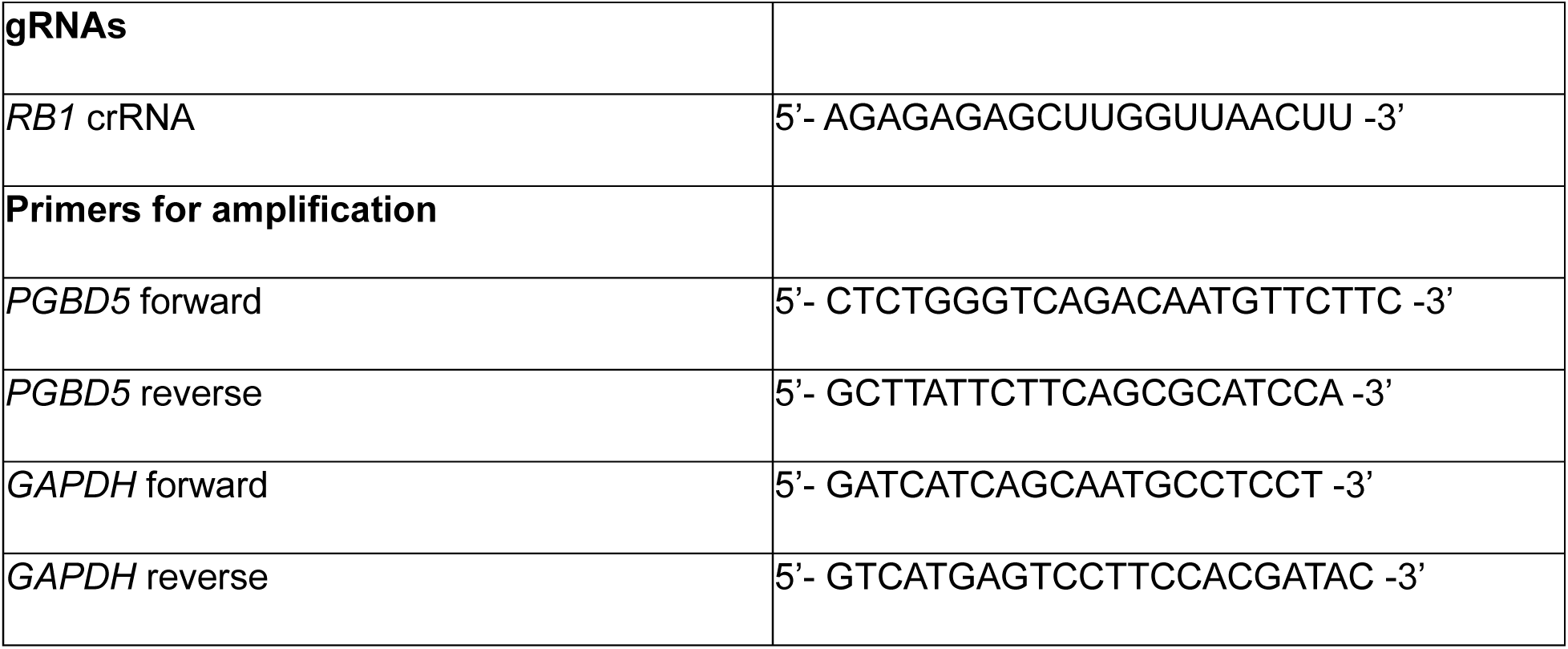
List of oligonucleotides.

### Western blotting

To assess protein expression by Western immunoblotting, pellets of 1 million cells were prepared and washed once in cold PBS. Cells were resuspended in 100-130 µL of RIPA lysis buffer (50 mM Tris-HCl, pH 8.0, 150 mM NaCl, 1.0% NP-40, 0.5% sodium deoxycholate, 0.1% sodium dodecyl sulfate) and incubated on ice for 10 minutes. Cell suspensions were then disrupted using a Covaris S220 adaptive focused sonicator for 5 minutes (peak incident power: 35W, duty factor: 10%, 200 cycles/burst) at 4 °C. Lysates were cleared by centrifugation at 18,000 g for 15 min at 4 °C. Protein concentration was assayed using the DC Protein Assay (Bio-Rad) and 15-35 µg whole cell extract was used per sample. Samples were boiled at 95 °C in Laemmli buffer (Bio-Rad) with 40 mM DTT and resolved using sodium dodecyl sulfate-polyacrylamide gel electrophoresis. Proteins were transferred to Immobilon FL PVDF membranes (Millipore), and membranes were blocked using Intercept Blocking buffer (Li-Cor). Primary antibodies used were: anti-EZH2 (Cell Signaling Technology, 5246) at 1:1,000, anti-RPA32 pT21 (abcam, ab109394) at 1:2,000, anti-RPA32 pS4/pS8 (ThermoFisher, A300-245A) at 1:2,000, anti-pCHK1 S296 (Cell Signaling Technology, 90178) at 1:250, anti-MYCN (Cell Signaling Technology, 9405) at 1:250, anti-Actin (Cell Signaling Technology, 4970 and 3700) at 1:5,000. Blotted membranes were visualized using goat secondary antibodies conjugated to IRDye 680RD or IRDye 800CW (Li-Cor, 926-68071 and 926-32210) at 1:15,000 and the Odyssey CLx fluorescence scanner, according to manufacturer’s instructions (Li-Cor). Image analysis was done using the Li-Cor Image Studio software (version 4).

### Transcriptomic data

Expression levels of *PGBD5* and *MYCN* in G401 cells were determined from our previously published dataset (8). Transcriptomic data and metaVIPER protein activity inference was performed as previously described (15, 16).

### Generating shPGBD5 cells

For shRNA cells, pLKO.1 shRNA vectors targeting *PGBD5* (TRCN0000138412, TRCN0000135121) and control shGFP were obtained from the RNAi Consortium (Broad Institute). Lentivirus production was carried out as described previously (8). G401 cells were transduced at an MOI ∼1.5 and selected with puromycin at 2 µg/mL for 72 hours. Knockdown was confirmed by quantitative RT-PCR as previously described (9), using primers specified in the Table 1.

### Combination drug treatment

Drugs used for *in vitro* treatment were supplied by Selleckchem (TAZ, S7128; Elimusertib, S9864; camptothecin, S1288; SRA737, S8253).

For combination treatment with TAZ and elimusertib, we used a two-dimensional dose matrix design, treating the cells for 9 days. After the addition of cells, drugs were added using a pin tool (stainless steel pins with 50 nL slots, V&P Scientific) mounted onto a liquid handling robot (CyBio Well vario, Analytik Jena). On Day 9, CellTiter-Glo reagent was freshly reconstituted and added in a 1:1 proportion to cell media, according to the manufacturer’s instructions. A similar protocol was used for monotherapy dose response curves for elimusertib and SRA737, although drug solutions were added manually, and specific treatment times are indicated in the corresponding figure legends. For analysis of synergy, we used the synergyFinder package (58). Outliers due to pinning errors were excluded after manual examination.

### Immunofluorescence

Immunofluorescence for γH2AX was performed on cells plated on Millicell EZ Slide glass slides (EMD Millipore), coated for 45 minutes with bovine plasma fibronectin (Millipore Sigma). After drug treatment, cells were washed once with PBS and fixed in 4% formaldehyde for 10 minutes at room temperature. Slides were then washed three times in PBS for 5 minutes, permeabilized for 15 minutes in 0.3% Triton X-100, washed again in PBS three times, and blocked with 5% goat serum (Millipore Sigma, G9023) in PBS for 1 hour at room temperature. Slides were incubated with mouse anti-γH2A.X primary antibody (Sigma-Aldrich, 05-636) at 1:500 in blocking buffer for 1 hour, washed three times in PBS, and incubated with goat anti-mouse secondary antibody conjugated to AlexaFluor555 (Invitrogen, A-21422) at 1:1,000. Cells were then counterstained with DAPI at 1:1,000 for 10 minutes and treated with ProLong Diamond Antifade Mountant with DAPI (Invitrogen, P36962) for 48 hours.

Images were acquired on a Zeiss LSM880 confocal microscope at 63x magnification. Images were then processed using a custom pipeline in CellProfiler (59). Per-cell integrated γH2A.X intensity was normalized against per-cell integrated DAPI intensity. All image analysis used the same pipeline settings, with the exception of the RescaleIntensity module for the AF555 channel, which used the settings 0.009-0.09 for the images in Figure 2 and 0.005-0.09 for Figure 4. CellProfiler analysis pipeline files and raw image files are available on Zenodo (DOI: 10.5281/zenodo.10982946). Overlaid images were prepared using Fiji (60).

### Xenografts

All mouse experiments were carried out in accordance with institutionally approved animal use protocols (8). To generate PDXs, tumor specimens were collected under approved IRB protocol 14-091, immediately minced and mixed (50:50) with Matrigel (Corning, New York, NY) and implanted subcutaneously in the flank of 6-8 weeks-old female NOD.Cg-*Prkdc^scid^ Il2rg^tm1Wjl^/Szj* (NSG) mice (Jackson Laboratory, Bar Harbor, ME), as described previously (61). Mice were monitored daily and PDX samples were serially transplanted three times before being deemed established. PDX tumor histology was confirmed by review of H&E slides and direct comparison to the corresponding patient tumor slides. PDX identity was further confirmed by MSK-IMPACT sequencing analysis.

Therapeutic studies used female and male NSG mice obtained from the Jackson Laboratory. Xenografts were prepared as single-cell suspensions, resuspended in Matrigel, and implanted subcutaneously into the right flank of 6-10 week old mice. 100 µL of tumor cell suspension was used for each mouse. Tumors were allowed to grow until they reached a volume of 100 mm^3^, at which point they were randomized into treatment groups without blinding. Drugs were prepared using the following formulations: Tazemetostat was dissolved at 25 mg/mL in 5% DMSO, 40% PEG 300, 5% Tween 80, and 50% water. Elimusertib was dissolved at 5 mg/mL in 10% DMSO, 40% PEG 300, 5% Tween 80, and 45% water using a sonicator. Drugs were reconstituted daily. TAZ was dosed at 250 mg/kg twice daily by oral gavage, 7 days per week. Elimusertib was dosed at 40 mg/kg twice daily by oral gavage using 2 days on and 12 days off cycle. Caliper tumor measurements were taken twice weekly. Tumor volumes were calculated using the formula Volume = (π/6) x length x width^2^. Tumor growth analysis was performed using the Vardi *U*-test (62), as implemented in the clinfun R package using the aucVardiTest function. Tumor-free survival analysis was calculated using OriginPro (Microcal) by the Kaplan-Meier method, using the log-rank test. Raw data and R scripts used for data analysis are available on Zenodo (DOI: 10.5281/zenodo.10398544).

---

**Supplementary Table S1:** List of PDX models used in this study, with clinical characteristics of the original tumor specimens, followed by a list of mutations found in all PDX models, as determined by targeted MSK-IMPACT sequencing.

