## Supplementary figures and images for "Epigenetic targeting of PGBD5-dependent DNA damage in SMARCB1-deficient sarcomas"

### Uncropped Blots

#### Slide 1
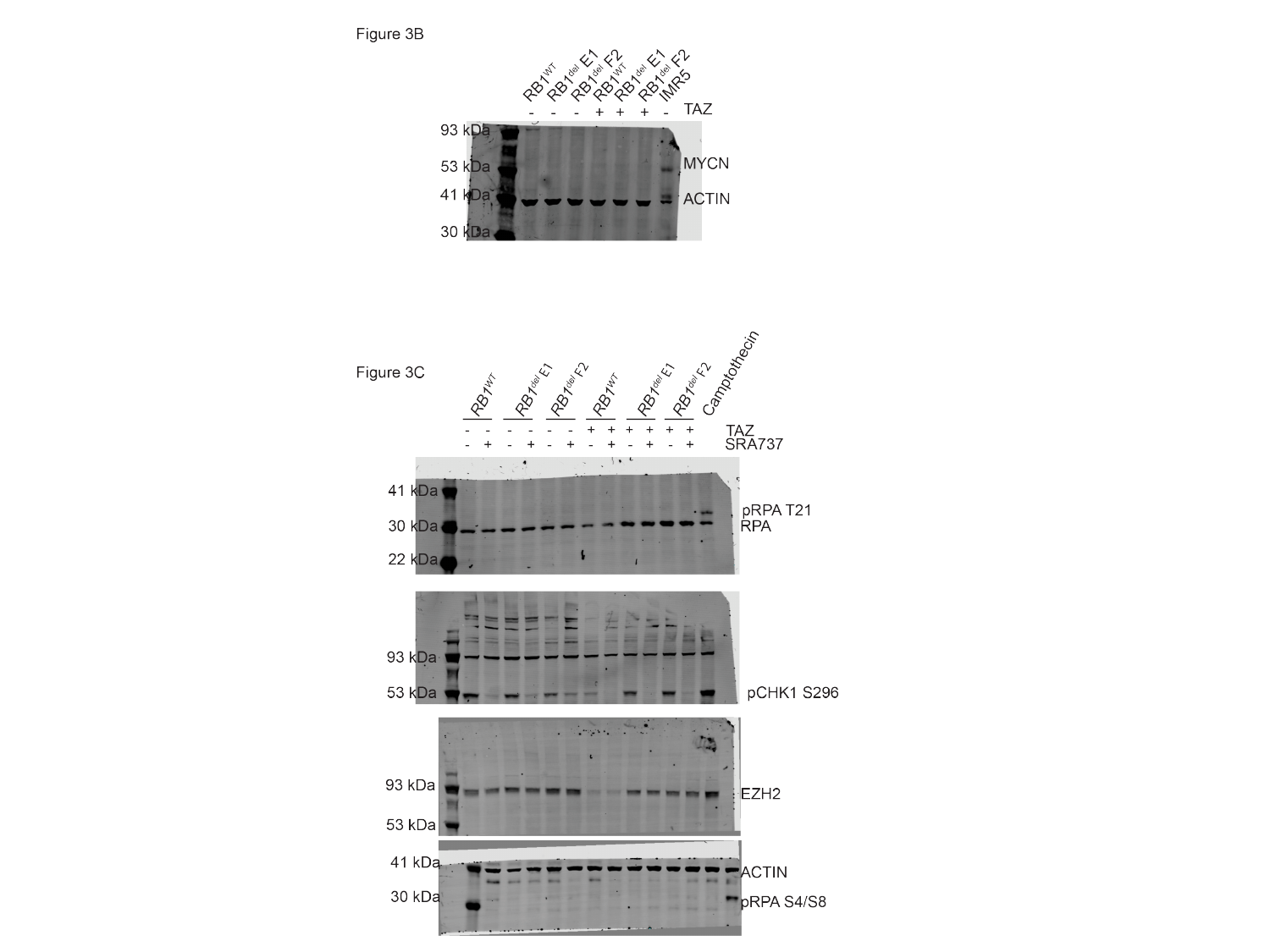
